# Analysis of Larval Fish Feeding Behavior under Naturalistic Conditions

**DOI:** 10.1101/2022.11.14.516417

**Authors:** Shir Bar, Liraz Levy, Shai Avidan, Roi Holzman

## Abstract

Modern research efforts concerned with animal behavior rely heavily on image and video analysis. While such data are now quick to obtain, extracting and analyzing complex behaviors under naturalistic conditions is still a major challenge, specifically when the behavior of interest is sporadic and rare. In this study, we present an end-to-end system for capturing, detecting and analyzing larval fish feeding behavior in unconstrained naturalistic environments. We first constructed a specialized system for imaging these tiny, fast-moving creatures and deployed it in large aquaculture rearing pools. We then designed an analysis pipeline using several action classification backbones, and compare their performance. A natural feature of the data was the extremely low prevalence of feeding events, leading to low sample sizes and highly imbalanced datasets despite extensive annotation efforts. Nevertheless, our pipeline successfully detected and classified the sparsely-occurring feeding behavior of fish larvae in a curated experimental setting from videos featuring multiple animals. We introduce three new annotated datasets of underwater videography, in a curated and an uncurated setting. As challenges related to data imbalance and expert’s annotation are common to the analysis of animal behavior under naturalistic conditions, we believe our findings can contribute to the growing field of computer vision for the study and understanding of animal behavior.

## 1 Introduction

Modern zoological research relies heavily on image and video acquisition to extract quantitative data on animals in the laboratory or in their natural environment. While video and image data are relatively quick to obtain, analysis capabilities remain a major bottleneck in extracting insights from the raw data [1–3]. This bottleneck is especially severe when the events of interest are sporadic, and are unpredictable in time and space. This is particularly true when sampling freely-behaving animals under naturalistic conditions. In this case, given a raw stream of data, such as a video, the behavioral events of interest make up only a very small portion of the data.

The scarce and highly variable data give rise to two challenges: first, it diminishes the statistical power for testing the biological question; second, it hinders the generation of large datasets that are required for the training and application of modern computer vision techniques based on Deep Learning [4].

Although many machine vision applications have been developed in recent years to track and monitor animals in a laboratory setting [reviewed by 5], completing this task in an unconfined, naturalistic environment with unknown number of animals is still an ongoing effort, especially in the aquatic realm. Specifically, many difficulties arise from the unpredictable nature of the animals and their environment. For example, how to image a large space; how to deal with visual challenges such as occlusions, animals going in-and-out of focus, and changing lighting conditions?

In this study, we introduce a novel system for imaging the feeding behavior of fish larvae under naturalistic conditions, and test it in a curated setting (see Fig. 1). The ultimate purpose of the system is to reduce the knowledge gap regarding the causes of the high mortality rates of larval fish reared in aquaculture, and to provide a tool for monitoring the well-being of larvae in these facilities. To attain this goal, we constructed an underwater camera system able to capture videos under naturalistic aquatic conditions. Two of the prominent features of this environment are (a) highly variable lighting, and (b) individuals that move in complex 3-dimensional tracks, often occluding one another or moving in and out of focus. To analyze these data, we utilize a detection and action classification approach. We localize rare behaviors such as feeding strikes (“strike”), as well as common behaviors (“swim”), in videos featuring multiple naturally-behaving animals.

**Fig. 1.**
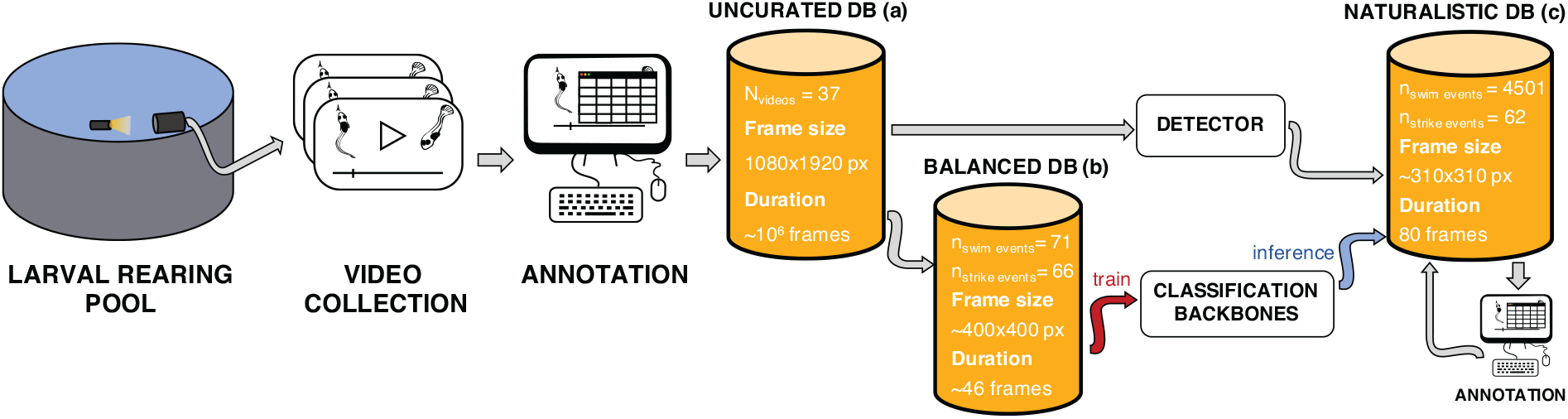
Overview of the study workflow. Starting from an aquaculture rearing pool on the left, our specialized system yielded high-speed videos of larval fish behavior. We annotated 29 million frames across 37 videos, searching for larval feeding strikes. These comprised the uncurated dataset (a). We cropped short clips around the most visually clear feeding events and, combining the with a similar number of clips of swimming larvae, we created the balanced curated dataset (b). This dataset was used to train our action classifier. To complete the pipeline, we trained a fish detection model, and combined the two models in a curated setting to detect and classify events from raw videos. In the process, we created and manually annotated the naturalistic curated dataset (c).

Using the data obtained from our system, we publish three new datasets of fish larvae behavior; in curated and uncurated settings. A single uncurated dataset consists in over 29 million frames, across 37 different video sequences; with all feeding strikes in each sequence manually annotated (Fig. 1a). In the curated setting, we present two datasets consisting in cropped clips: a balanced dataset with the most visually coherent “strike” samples from all our videos, i.e., minimal occlusions and reduced blurriness; and a balanced class structure for both “swim” and “strike” behaviors (Fig. 1b). In addition, we present a more challenging naturalistic dataset, with over 4,500 cropped clips, posing challenges both in terms of imaging conditions (occlusions, high degree of blur, multiple individuals) and in terms of a highly unbalanced class structure, where the class of interest is rare (Fig. 1c). All three datasets were acquired using two camera setups, several different illumination settings, and several different filming arenas (i.e., fish rearing tanks).

We discuss the various challenges of our work in the following sections. We address the challenges of deploying an underwater camera system in section 3.2, and the labeling efforts required to collect and annotate a large corpus of fish swim/strike behaviors in section 3.3. The design of computer vision algorithms for the detection and classification of fish actions is discussed in sections 3.4 and 3.5, and their results are reported in section 4. Finally, we compare the performance of our two top models and discuss sources of error and implication to the deployment of the system in the field in section 4.3. These challenges are common to the analysis of animal behaviour under naturalistic conditions, and our findings and solutions contribute to the growing field of computer vision for animals.

## 2 Related work

In this section we acquaint the reader with some of the relevant background. We start by addressing the biological problem in regard to larval fish feeding (section 2.1); describe previous work and the challenges of underwater photography (section 2.2); followed by a short review of previous work in action recognition, pertaining mostly to the field of human behavior (section 2.3). We then specify several different approaches in modeling animal behavior, with emphasis on the particular challenges of imaging and analyzing fish behavior (section 2.4).

### 2.1 Fish larvae feeding behavior

Most marine fish reproduce by releasing fertilized eggs into the open ocean. The hatching larvae are undeveloped compared to the adults, and metamorphose to obtain the adult morphology and characteristics within a period of several weeks.

During this period, the larvae experience prodigious mortality rates. In the ocean, more than 90% of the larvae will die within the first 30 days post-hatching [6].

This mortality rate has been attributed to various mechanisms, including the inability to find food, and death from predation or disease [7]. However, even in aquaculture rearing pools, where conditions are supposedly ideal, the mortality rate remains above 70% [8]. Previous imaging efforts in the laboratory implicated larval inability to successfully capture their prey as the major instigator for hunger and starvation-induced mortality [9, 10].

Critically, the feeding behavior of larval fish has never been imaged outside the confines of the laboratory. Such capacity would be key to understanding how larval feeding is influenced by a complex array of environmental conditions. Our inability to quantify this extremely rare behavior outside the laboratory is partly due to the technical challenges involved in underwater videography (see below), but also due to the highly sporadic nature and rarity of feeding events initiated by the larvae.

### 2.2 Underwater computational photography

Underwater camera systems must deal with the challenges presented by the surrounding medium (i.e., water). The main technical challenge of underwater imaging lies in the limited viewing distances of objects, stemming from the effects of turbidity and light attenuation, which limit the coverage of a habitat compared to filming in air [11, 12]. On land, medium sized animals such as birds can be viewed at distances of 100s of meters [13] while even in the clear waters of coral reefs, similar-sized fish can only be viewed from few 10s of meters at best [14, 15]. Underwater images also tend to flicker due to variations created by the refraction of sunlight through the dynamic undulating air-water surface, and a method to correct this, in the case of stereo cameras, was proposed by [16]. In addition, recovering scene geometry in the case of light traveling through air and water is a challenging task, and a relevant theory to resolve this was posited by [17].

### 2.3 Action recognition

Action recognition has been developed, by and large, with a focus on human behavior. Recognizing actions or behaviors of subjects in video data can be divided into detecting the action in a spatio-temporal sequence, and classifying the action. For action classification, there have been several successful variations of 3D Convolutional Networks (3D-ConvNets) in recent years - Two-Stream [18], I3D [19], P3D [20], SlowFast [21].

Notably, the Two-Stream and the two-stream I3D models utilized optical flow as an additional input stream into the network to capitalize on fine-grained motion data.

SlowFast [21], also employs a two-stream architecture. However, rather than using pre-computed optical flow, SlowFast varies the sampling rate of the input video in each of the streams in order to facilitate the learning of different features. The two streams are homologous except for their channel depth and sampling rates. The Slow pathway samples at a lower frequency but has a deeper structure, aimed at capturing spatial features. The Fast pathway samples more frames but has fewer channels in every block, aimed at targeting motion features.

In addition to action classification, SlowFast also has an action detection module, which is akin to Fast-R-CNN [22]. The module uses a pretrained human detector based on a Faster R-CNN [23, 24], to generate region proposals and was trained on 15-minute-long videos from the Atomic Video Actions (AVA) dataset [25].

### 2.4 Animal behavior modeling

Fish are particularly challenging to model and track. As they move in a 3D volume, their projected silhouette changes frequently. Additionally, as fish tend to school, individuals often cross trajectories and occlude one another [26]. Typically, a considerable amount of work is necessary to generate even a modest dataset of fish behaviors, even under controlled conditions. For example, [26] created a dataset of 3D-zebrafish tracking in a laboratory setup using a tracking-by-detection methodology, comprising 8 sequences, each between 15-120 seconds long.

A popular approach to the analysis of complex behaviors from video sequences is that of markerless pose estimation [e.g., 27, 28]. Using this approach, rather than applying action/behavior classification to the raw videos, the positions of body parts are extracted using pose estimation, and the analysis of behavior happens on a time-series of marker locations. Pose estimation alone, however, does not provide behavior classification, and additional algorithms are required in order to translate marker sequences into behaviors [5].

In general, using action classifiers to classify animal behaviors directly from video is not common. Long et al., [29], studied nest-building behavior in Cichlid fish in a laboratory setting using a combination of classic vision methods and a variant of 3D-ResNet action classifier [20]. They used Hidden Markov Models (HMMs) to detect disturbances in artificially-colored sand as a proxy for the location of behaving fish. While their approach produced a large dataset of fish behaviors, the detection required a specific, manipulated environment (i.e., artificially-colored sand) that is difficult to reproduce outside of the laboratory. In addition, the behavior of interest only occurred along the 2-D bottom, bypassing challenges associated with 3-D complex behaviors.

Only few attempts have been made to automate the analysis of fish behavior from videos under naturalistic conditions. For example, [30] used tracking-by-detection obtain the 3D trajectories of free-swimming reef fish *in situ* using relatively long video sequences. However, their system was only semi-automatic as all their trajectories were corrected by means of human in the loop.

Shamur et al., [31], attempted to classify the feeding behavior of fish larvae in a laboratory using classic vision methods. In their work, larvae were detected and classified using a pipeline of edge detection, action descriptors extractions, and SVM classifiers. Although they achieved reasonable results (AuROC = 0.82, ACC = 72.7±2.1), the analysis pipeline is not transferable to naturalistic settings, such as ours. Their pipeline requires manual tuning of thresholds for each new video analyzed to adjust for changes in lighting conditions and water clarity; the edge detection function works poorly for out-of focus fish, which are common in our data; Image quality in the laboratory was superb compared to our *in-situ* filming because in the laboratory the water are cleaner and most of the optical path is traversed through air. For this reason, clips in the laboratory dataset were cropped around the mouth of the fish, not visualizing the whole body, unlike our dataset. In addition, to avoid occlusions and maintain fish in the focal field, the larvae were placed in a narrow arena (∼ 5mm), essentially creating a 2D scene. As a by-product, this confined space potentially limited the natural behavioral repertoire of the fish. For all these reasons, we do not assess our models on this dataset, nor run the old pipeline on our new data.

## 3 Methods

In this section, we present the development a unique analysis pipeline for the detection of larval fish behavior in aquaculture rearing pools, from the initial infrastructure to the deep learning models.

We first briefly describe the study system and discuss the particular challenges and the novelty of building an imaging system to visualize such tiny, fast-moving creatures underwater (sections and 3.2). We then discuss the intensive effort of collecting and annotating the data produced by this system into one uncurated and two curated datasets (section 3.3). Next, we discuss the main powerhouse of our analysis pipeline; the classification module and its training, the specific challenges of training on very low sample sizes, and how we chose to tackle them (section 3.4). Finally, we explain how we combined our classifier with a fish detector to create an analysis pipeline and tested it in a curated fashion (section 3.5).

### 3.1 Study system

The feeding behavior of larval sea-bream was recorded over the course of 17 months in several rearing tanks located in a commercial aquaculture facility (ARDAG hatchery, Eilat, Israel). We deployed a submerged camera system in the tanks to record the feeding in freely-behaving larval fish, from age 7 DPH (days post hatch) to 32 DPH. In this facility, larvae are reared in large circular tanks (∼ 4 m in diameter and ∼ 1.5 m deep, Fig. 2a) under controlled temperature and oxygen conditions. Cohorts of ∼ 10^6^ eggs are introduced into each tank, where they hatch and are grown until they metamorphose to adult-like morphology at ∼35 DPH. Larvae are fed twice a day with various food types (Rotifers, Artemia, and pellets) according to their age and size.

**Fig. 2.**
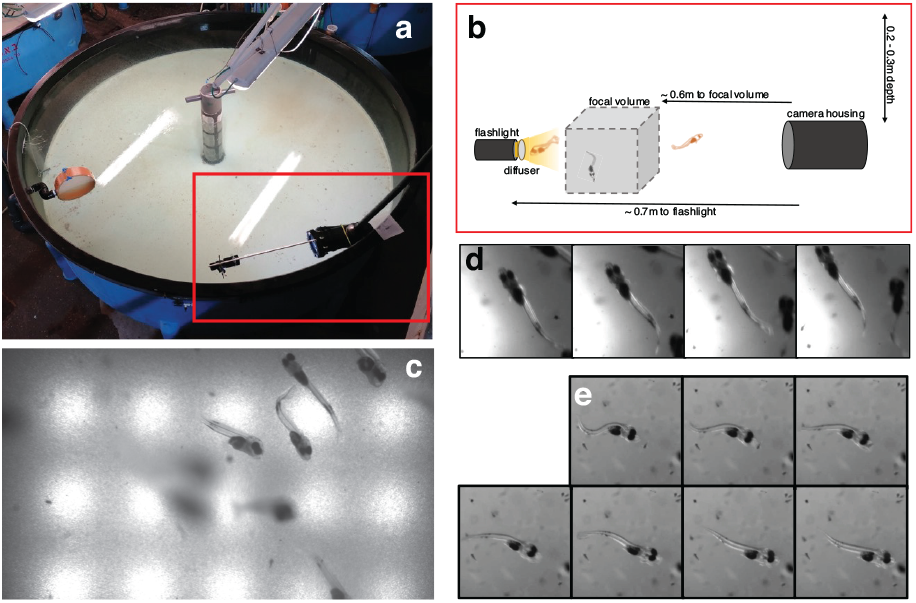
The camera system and data acquisition pipeline. a) the camera system in a fish larvae rearing pool at the ARDAG hatchery, Eilat; b) a schematic depiction of the submerged camera setup, including a representation of the focal volume, where fish appear sharpest; note that this volume is a mere 0.002 % of the pool c) an example of a full video frame acquired using the system. Larvae appear in various levels of sharpness within the frame, see Fig. S1 in Supplementary Information (SI) for more examples; d) an example of a “swim” sequence from the naturalistic dataset; the uneven lighting and multiple fish are typical of this dataset, see clip (Online Resource 1); and e) a representative sequence of a “strike” event from the balanced dataset, note the typical S-shape assumed by the fish and the even lighting typical of balanced dataset, see clip (Online Resource 2). Panels (a,b) pertain to the first camera setup.

Larval fish (like most of the adults) capture their prey using suction feeding. In this feeding mode, the fish abruptly open the mouth and expand the buccal cavity to generate an inflow of water into the mouth that carries the prey in with it. Suction-feeding is an extremely rapid behavior, with events typically taking less than 40 milliseconds from the start of mouth opening to mouth closing. Larval fish are minuscule, hatching at a body length of ∼ 3 mm, and growing to ~10 mm at 30 DPH. Correspondingly, both their mouth and prey diameter measure ca. 0.1-1 mm. The size of the animals, their speed, and erratic 3D motion therefore necessitated the development of a specialized filming system.

### 3.2 Camera setup

To record the dynamics of the abrupt feeding events by the tiny larvae at an appropriate resolution, we needed to use a high-speed camera equipped with an ultra-zoom lens. We chose to use a submerged setup both because previous work has shown that feeding events are best characterized from a side view [10, 31, 32], and because the opaque walls and floor of the tank did not allow imaging from outside it or from above.

Filming was done using a high-speed monochrome camera (Optronis CP70-2-M/C-1000) connected to a computer and able to record relatively long videos (20-30 min) at a high frame rate of 750 frames per seconds (fps) and a resolution of 1920 *×* 1080 pixels. The camera was enclosed in an underwater housing and connected through four 30m long camera-link cables to a nearby computer (Fig. 2 a,b). The camera was equipped with a Navitar 6000 ultra-zoom lens providing 2:1 magnification (i.e., 2 mm in the real world are mapped on 1 mm of the camera’s sensor) with a large depth-of-field of 50 mm.

Overall, the setup enabled visualizing events within a volume of 40 × 60 × 50 mm (*H* ×*W* × *D*). The fast frame rate of the camera, water turbidity, and the lens’s limited aperture (f=8) required strong illumination, which was provided by a battery-operated SCUBA flashlight (Scubatec US15 LED 10 Watt). The flashlight was equipped with a diffuser and was positioned directly in front of the camera, providing backlight illumination. Light intensity was approximately equivalent to that of sunlight at 5m depth in clear coastal water [33]. The system was submerged to ∼ 0.2-0.3 m depth, and was removed from the tank after each deployment. The flashlight was rigged to the setup with a flexible detachable arm. As a result, the position of the flashlight varied between deployments; creating variable lighting conditions even between videos taken in the same tank on consecutive days using the same setup.

Two distinct camera setups were tested. The change in setup stemmed from an attempt to improve the sharpness of the videos. Initially, the camera was placed in a snug housing, leaving ∼ 0.6 m of water between the lens and the focal plane. Because the water in the rearing tanks is turbid, the resulting images were relatively blurry. We attempted to improve the setup by placing the camera in a longer housing, such that most of optical path was through air (inside the housing), rather than through water. This resulted in much sharper images with larger areas appearing in focus, but the edges of the housing were also sometimes included in the frame. It is important to note that even though both setups had their shortcomings, feeding events were still detectable by our analysts and the videos were successfully annotated.

Unlike the previously published dataset [31], which was acquired in the laboratory under constant conditions, videos in our dataset vary along many different axes. In addition to the visual differences caused by the two camera setups, videos varied considerably within each setup. This is because our filming documented the variable conditions within the rearing tank, as determined by the hatchery’s rearing protocols. Specifically, the age of the larvae, the number of larvae in the tank, the type and amount of food, water turbidity, illumination, *O*_2_, currents and turbulence, all varied between filming days; affecting both larval behavior (e.g. feeding rate) and the visual appearance of the videos (see Fig. S1 in S.I.).

### 3.3 Data curation

Overall, we obtained and annotated over 29 million frames across 37 videos, taken in different rearing pools over the course of 17 months. To manually annotate the (sparse) feeding events, a trained observer watched the video at 15 FPS and noted the time and coordinates of all feeding-related events. Often, the observer had to re-play the video to ascertain an observation. We estimate that annotating our datasets took a minimum of 540 work hours. This is an underestimation as we are not allowing for data entry or possible re-plays. Feeding events, or “strikes”, were defined as events that started with the larva assuming an S-shape position, followed by a rapid forward lunge and opening of the mouth (Fig. 2e). These events are visually distinct and represent high-effort prey-acquisition attempts that are likely to be successful [10]. Note that this definition is different than the one used by [31], there filming conditions allowed for a high resolution visualization of the fish’s mouth, which was not possible under the conditions of the rearing pools.

#### 3.3.1 Uncurated dataset

Our uncurated dataset is the set of all raw videos collected, 29 million frames in 37 videos. The annotation provided is the temporal and spatial coordinates of all feeding strikes, after an exhaustive annotation effort which yielded a total of 90 events. Of the videos, 21 were from the first setup (8, 436, 822 frames) and 16 from the second setup (20, 805, 438 frames). Each video comprised a monochrome sequence, with mean length of 790,331 frames.

#### 3.3.2 Balanced Curated dataset

From the uncurated corpus we create a balanced on which we trained our classifier.

All 90 “strike” events were manually inspected to exclude samples in which the fish appeared severely blurred, occluded, or in low-contrast (thus removing *n* = 24 events). For each of the remaining 66 visually coherent strikes, we extracted a spatially cropped square clip around the larva of interest. Unlike [31] who cropped clips centered around the larva’s mouth, our clips contained the entire body of the larva. This allowed us to incorporate pre-strike behaviors such as aiming at prey, which involve the whole body. We also temporally cropped each clip, starting 10 frames before the mouth opened and ending 5 frames after the mouth closed. These cropping procedures resulted in a distribution of clip lengths and sizes (average±Sd duration= 45±18 frames; average±Sd height= 385±93 pixels).

For the “swim” events, we used a methodology based on Canny edge detection [34] to automatically detect potential larvae within a frame. Around each of these detections, we created cropped square clips, each 200 frames in length. Research assistants filtered out clips without larvae and spatially cropped each video as tightly as possible around the larvae. To avoid biasing the dataset due to the difference in clip duration between the “swim” and “strike” event classes, we further temporally cropped “swim” clips at random, in order to match the distribution of clip duration in the “strike” class.

From the available “swim” clips we selected 71 clips to obtain a balanced dataset design between the two action classes. Although it is a common practice to train the classifier on an event distribution similar to the one expected to be found in production (i.e., in the natural environment), our preliminary attempts to train a classifier on an unbalanced dataset produced poor results. This was probably due to the small size of the positive class, which we could not increase.

Our balanced dataset therefore comprised 66 clips featuring “strike” events, and 71 “swim” events that were further partitioned to train, validation, and test datasets. We ensured that clips cropped from the same raw video were grouped in the same partition in order to avoid data leakage. Additionally, to avoid creating spurious features related to differences in filming setup between videos, we selected a similar proportion of clips from each filming setup in each of the two classes. The training dataset comprised 41 “swim” and 39 “strike” clips; the validation set comprised 11 “swim” and 11 “strike” clips; and the testing set comprised 19 “swim” and 16 “strike” clips.

### 3.4 Classification module

Our pipeline consists in two models, trained separately: an action classifier and a fish detector. The action classifier was trained on the curated balanced dataset (section 3.3.2), and its performance was further evaluated on the naturalistic curated dataset, created using the detector. The following section describes the classification module.

#### 3.4.1 Model

We compared the performance of several popular backbones for the action classification of “strike” and “swim” events: an I3D network [19]; a two-stream SlowFast Network [21] with a 3D-ResNet-50 [35] backbone; and just the Slow pathway of the SlowFast network.

In the SlowFast model, the rate at which each pathway samples frames from the input clip is a user-specified hyperparameter. We chose to follow one of the settings suggested in the original paper [21], with the Slow pathway sampling eight frames uniformly throughout the clip, and the Fast pathway sampling 32 frames throughout the clip. To ensure that this sampling provided good coverage of the feeding strikes, the sampled clips were manually inspected. The ratio between the channels in each pathway, specified by the *β* parameter, was set to *β* = 1/8, as we used pre-trained weights (see below). Specifications of the rest of the training hyperparameters are provided in the Supplementary Information (SI, section S2.1).

#### 3.4.2 Training under a low data regime

Our chief challenge with training this module was the limited size of the balanced curated dataset, which contained fewer than 100 samples per class. We tackled this data scarcity using three complementary approaches: Transfer Learning [36]; intensive data augmentations tailored for our dataset; and variance image calculation.

Rather than training our models from scratch, we used Transfer Learning to fine-tune existing model weights. All three backbones we compared (I3D, Slow and SlowFast) were pre-trained on the Kinetics-400 dataset [19] - a dataset of 400 human action classes, with over 400 clips per class.

To asses the contribution of pre-training to model performance, we trained a SlowFast network from scratch on our dataset. We also tested whether using a model pre-trained on a different dataset might improve performance. For this, we use a SlowFast network pre-trained on the Something-SomethingV2 dataset (SSv2) [37]. We were inspired by works such as [27] which showed that pre-training on human pose data is beneficial when learning animal pose, in spite of the difference in domains. We chose the SSv2 dataset because recent work suggested it encourages the learning of more dynamic, temporal-related features [38]. In all experiments, we used the publicly available models and weights in the PyTorchVideo repository [39, 40]. We fine-tune all models for 50 epochs, full details are provided in section S2 in the SI.

Our dataset, though small, showed a diversity of visual conditions; mainly differences in lighting intensity and degree of blurriness of the fish (see Fig. S1 in SI). To encourage model generalization over these conditions and to enhance the number of samples in our dataset, we randomly applied augmentation to the intensity values of clips, varying the degree of brightness, and augmented the sharpness of clips by randomly applying Gaussian blur to samples during training.

We integrated our knowledge of the biology and behavior of the fish to generate an additional channel of information in our clips. We exploited the fact that “strike” behavior is characterized by abrupt movements, while “swim” behavior is typically a smooth undulatory movement. These differences are expected to affect the rate at which pixels change their intensity values throughout the clip, with “strike” pixels showing areas of higher variance.

Rather than calculating the optical flow for each clip, which is computationally intensive, we calculated the variance image of the entire clip (see Fig. S2 in SI). The variance image was duplicated along the temporal axis and stored as a third channel, alongside two duplicate channels of the clip’s monochrome sequence. We tested the contribution of this manipulation with an ablation study (section S2.4 in SI).

We note that given a larger dataset, we would expect our classification module to learn such features independently, making this additional channel superfluous. However, in light of our low data regime, we considered that this manipulation would inform the classifier and enhance learning.

### 3.5 Detection module and the entire pipeline

Equipped with the classifier trained on the balanced curated dataset, we moved towards a pipeline for analysis of longer videos rather than cropped clips. The pipeline comprised a detection module, followed by the classifier, as discussed next.

For the detection module we trained a Faster-RCNN [23] object detector with a ResNet-50-FPN backbone [24, 41] using the Detectron2 framework [42]. This detector was pre-trained on ImageNet [43] and fine-tuned on our detection dataset. Information on dataset, training procedures, and results are provided in SI (section S3).

After training our detector, we used it in conjunction with our behavior classifier (section 3.4) to create the behavior detection pipeline (Fig. 3) and generate the naturalistic dataset. To test our pipeline, we used 11 videos from the uncurated dataset that featured the highest number of feeding strikes according to the manual annotation. Rather than going through the entire video, we applied the pipeline on randomly sampled frames. In addition, we strategically sampled frames in the temporal coordinates of our manually annotated feeding strikes. In total, we sampled 62 strike-related frames and 949 frames at random. We maintained a ratio of ∼ 1:15 between frames that contained events of interest and those that did not, to emulate the natural rarity of larval feeding strikes.

**Fig. 3.**
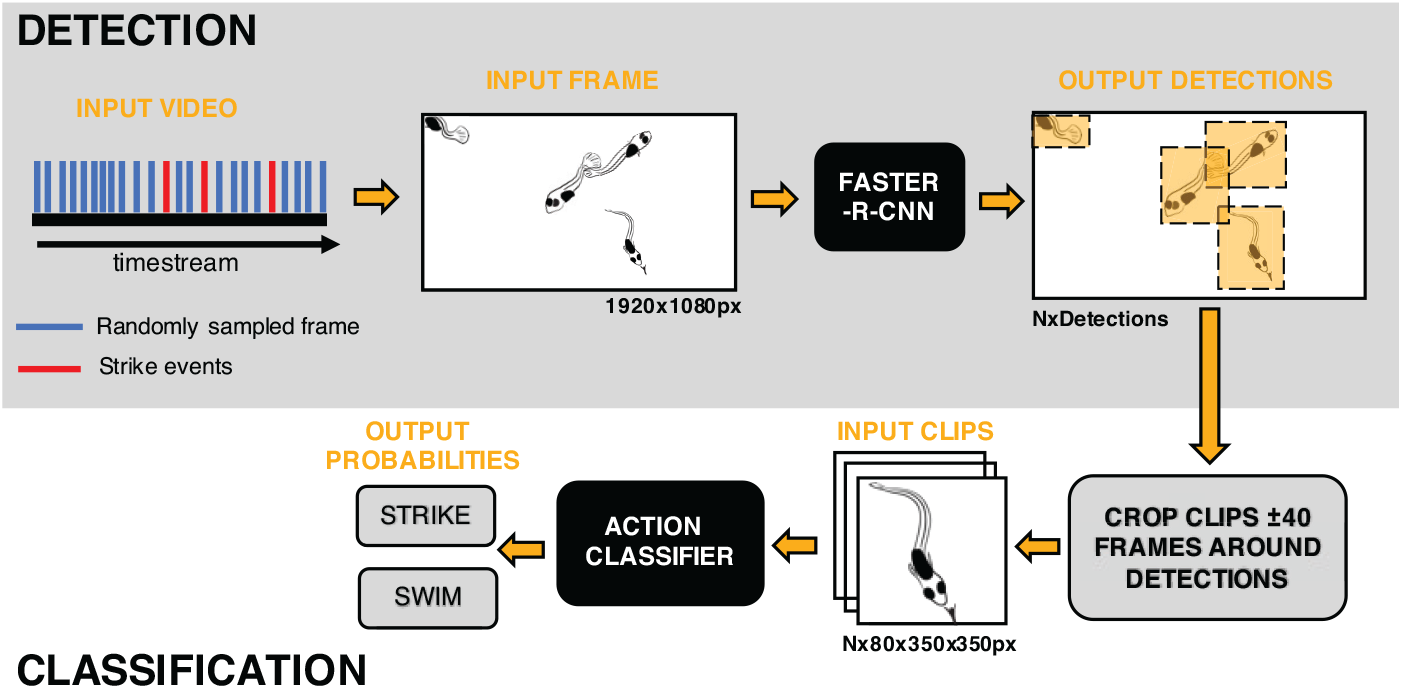
A diagram of the analysis pipeline and the generation of the naturalistic curated dataset. The flow depicted in this diagram, starting in the upper left corner, shows how we applied our analysis pipeline to frames sampled from long videos sequences, and generated clips of behaving fish to feed to the action classifiers. Using this scheme we created a naturalistic dataset of *n* = 4,563 clips which was further manually annotated.

For each frame sampled, the detector was applied to locate the fish in the frame (see Fig. 3). Around each of these detections, we created short cropped clips centered around the putative fish. Temporally, a ±40 frame window was cropped around the detection frame. Spatially, small square clips were cropped around each detection, with the cropping size determined according to the typical size of the fish in each video (range: 250-650 pixels). The variance image of each clip was calculated and the manipulated input was fed into the action classifier in order to obtain classification scores for each clip. As the class of interest was that of feeding strikes, from here on we refer only to classification scores relating to the “strike” class, hereinafter, “strike scores”. We provide additional technical details on implementation of the pipeline in section S4 in the SI.

There was a partial overlap between raw videos used in the balanced dataset and those used in the naturalistic dataset. Five of the 11 videos were used in the naturalistic dataset and in the test set of the balanced dataset. Six out of the 11 videos used in this dataset were also used in the train or validation splits of the balanced dataset. While this overlap is not optimal, we note that the clips in each dataset were generated differently. Temporally, samples in the naturalistic dataset are not tightly cropped around the feeding event, and actually clips are twice as long as an average feeding event; spatially, the clips were not tightly cropped around the fish of interest. Furthermore, four of these 6 videos featured additional strike events that were not included in the balanced dataset (*n* = 13 clips, range: 2 − 5 per video). When evaluating our classifier on this dataset we report results for the entire corpus of data, as well as for only those clips from the five videos not included in the training split of the balanced dataset, which we term the naive set.

### 3.6 Metrics

To evaluate the performance of our classification module on the balanced and the naturalistic dataset, we cast the problem as a binary classification problem, with “strike” being the positive class and “swim” being the negative class.

As customary in binary classification evaluation, we use the Receiver Operator Curve (ROC) [44], to obtain an estimate of the overall classifier performance across the entire range of decision thresholds. The area under this curve (AuROC) is often used to give a single score to the quality of the classifier. As our naturalistic curated dataset was highly imbalanced, we further evaluated the classifier using the Precision Recall Curve (PRC), following best practice delineated by [45], and used the area under this curve (AuPRC) as an additional quality score for both datasets. As noted by [46], PRC is sensitive to the class imbalance of the dataset, with expected curves changing according to the positive class percent in the data, and classifiers have been shown to produce drastically different results under changing data imbalances [46]. For this reason, our reported AuPRC are true for the stated class imbalance. All metrics were calculated using the precrec package in R [47].

## 4 Evaluation/Experimental Results

In the following section, we report the results of our classifier training and the performance of the entire analysis pipeline. We compare the performance of the three different backbones on our naturalistic dataset and asses the contribution of pre-training to classifier’s performance. We also present an analysis of the possible sources of error in our pipeline’s performance, and the insights gained from the manual annotations of this naturalistic curated dataset.

### 4.1 Classifier balanced curated results

Performance of all of the backbones on the balanced dataset is described in Table S2 and Fig S3. All backbones achieve decent results on the balanced dataset, except the SlowFast network with no pre-training that was not better than a random classifier. The SlowFast pre-trained backbones were the best-performing, with the SSv2 being superior to the Kinetics by a small margin. Both networks attained an AuROC of 1 on the train split. For the validation split, the AuROCs were 0.85 and 0.98 for the Kinetics and SSv2 pretrained SlowFast networks, respectively. For the test split, the AuROCs were 1 for both. Note that our best models achieve better AuROC scores than those obtained previously by [31] on their laboratory dataset.

### 4.2 Pipeline performance and the naturalistic curated dataset

The combination of the detection and classification modules yielded the naturalistic curated dataset (as depicted in Fig. 3). This dataset poses challenges typical of naturalistic conditions, such as multiple occluding animals, extreme lighting conditions, motion blur, and severe class imbalance.

The resulting dataset comprises 4,563 short clips (62 “strikes” and 4,501 “swims”); with the positive class accounting for ∼ 1.4% of the data. The entire dataset was annotated by a team of trained observers using a custom labeling software. To prevent bias, all observers were blind to the results of the classifier. For each clip, observers labeled the main activity of the fish in the clip. Observers also noted parameters regarding the behavior of the fish and the photographic quality of each clip (see Fig. S7 in SI). We merged these behavioral and photographic labels into the following mutually exclusive categories: strikes, abrupt movements, non-routine swimming, compromised footage, routine swimming, and can’t tell or no fish (see section S5.3 in SI for details)

Results of all backbones on the dataset (n=4,563), are given in Table 1 and Fig. 4. As with the balanced dataset, the pre-trained Slow-Fast backbones showed superior performance with a mean AuROC of 0.94, 0.97 and mean AuPRC of 0.6, 0.66 for the Kinetics and SSv2 respectively. Note that the expected AuPRC for a random classifier under the observed class imbalance is 0.014. The I3D and Slow backbones performed considerably worse than on the balanced dataset. The model with no pre-training performed poorly, being in line with a random classifier. Results for the naive video set, excluding videos used for the train split of the classifier, were similar (Fig. S5 in SI).

**Table 1.**
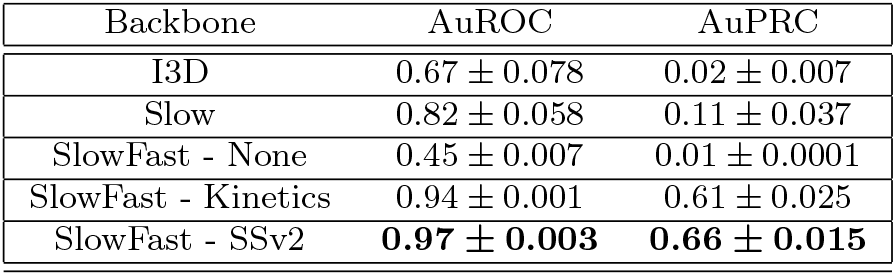
Model performance on the naturalistic dataset. Numbers represent *meanAuC ± std* for each backbone, averaging results from training using three different random seeds, I3D and Slow backbones were pre-trained on Kinetics. The pre-trained SlowFast variants are by far the best performing in both metrics.

| Backbone | AuROC | AuPRC |
| --- | --- | --- |
| I3D | $0.67 \pm 0.078$ | $0.02 \pm 0.007$ |
| Slow | $0.82 \pm 0.058$ | $0.11 \pm 0.037$ |
| SlowFast - None | $0.45 \pm 0.007$ | $0.01 \pm 0.0001$ |
| SlowFast - Kinetics | $0.94 \pm 0.001$ | $0.61 \pm 0.025$ |
| SlowFast - SSv2 | **$0.97 \pm 0.003$** | **$0.66 \pm 0.015$** |

**Fig. 4.**
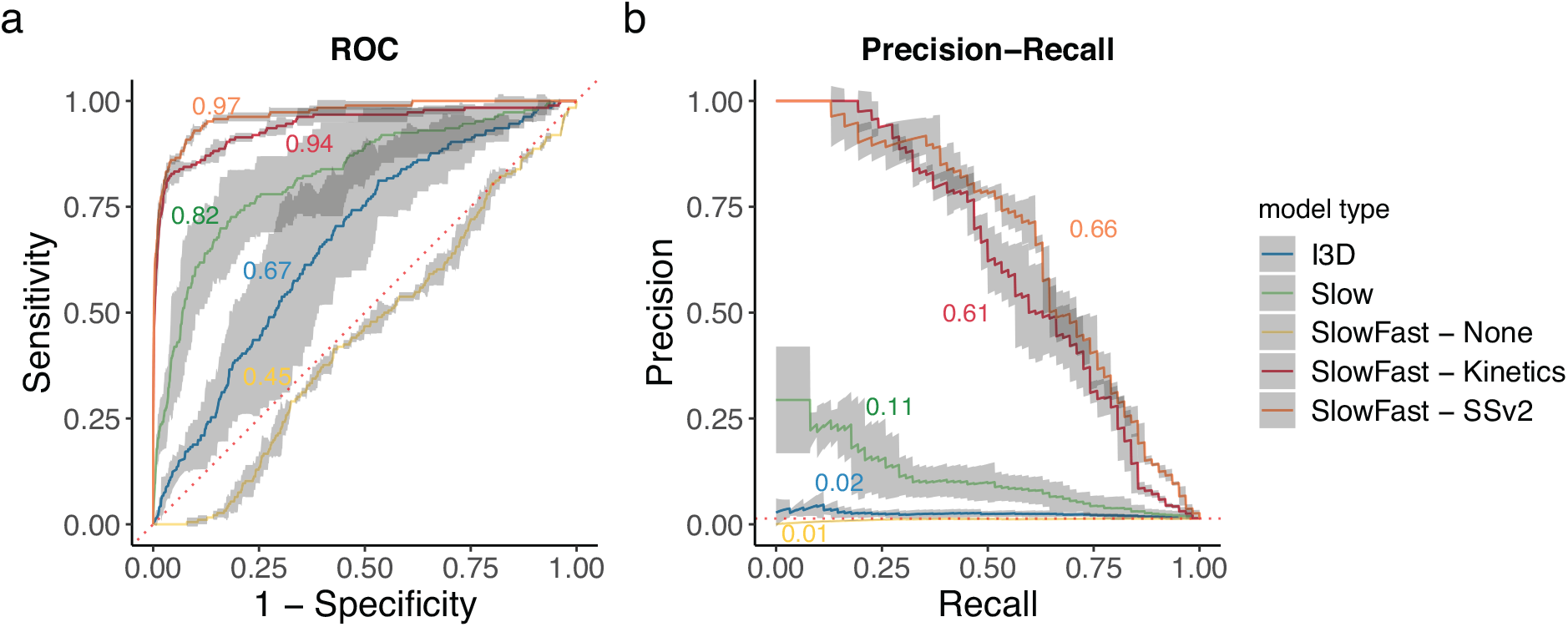
Comparison of backbones on the naturalistic curated dataset. a) ROC, plotting the True Positive Rate (Recall/Sensitivity) against the of False Positive Rate (FPR/1-Specificity); b) PRC, presenting the Precision as a function of Recall. In both a & b dashed red lines represent the performance expected by at random classifier. Each curve is the mean ROC and PRC for each backbone, averaging performance on 3 different random seeds. Shaded area around each curve are the 95% confidence bounds. The I3D and Slow backbones (blue and green respectively) are pre-trained on Kinetics, the SlowFast are either not pre-trained (yellow), or pre-trained on Kinetics (red) or SSv2 (orange). The two pre-trained SlowFast backbones (Kinetics, SSv2) show superior performance by a large margin.

These results show, that despite the extremely low data regime, our training resulted in models that are capable of detecting larval fish feeding behavior under challenging naturalistic conditions.

### 4.3 Error analysis

We investigated the possible sources of classification errors by analyzing the human annotations of the curated naturalistic dataset (N=4,563, see section 4.2) and the strike scores assigned to them by the two best models (SlowFast pre-trained on Kinetics and SSv2). We analyzed two rather than just one best model as the two present an interesting performance trade-off between the action classes (see below).

According to the human annotation, the detector had misclassified 118 samples as fish (∼2.5%), where no fish were found. Approximately 50% of the clips were labelled as high-quality clips featuring routinely swimming fish (n=1,841). Other distinct larval behaviors were those of abrupt movements (n=348) and non-routine swims (n=498; interrupted and reverse swimming, floating). The data also included a high percentage of low quality clips with compromised footage (n=1,196), featuring over-exposed imagery, a moving background induced by strong flows, or extremely blurred fish. The SSv2 pre-trained SlowFast performed consistently better than the Kinetics one in both AuROC and AuPRC scores (see Table 1). However, when examining the the distribution of strike scores per class (Fig. 5), it transpired that the improved performance of the SSv2 variant comes from a better mapping of the swim class to low strike scores (leading to higher rate of true negatives), but this comes at a cost of missing ∼ 16% of the strike events (more false negatives). Conversely, the Kinetics variant is worse at mapping the swim class, with more potential false positive detections, but better at the strike class, with only 1-2 events being mapped to low strike scores (more true positives). Figure 5 shows that the kinetics variant was strongly affected by abrupt movements and low-quality clips, which were disproportionately mapped to higher strike scores > 0.5, potentially explaining some of the classification errors. The SSv2 variant assigned low strike scores to compromised footage, but also gave disproportionately high scores to abrupt movements (Fig. 5 inset).

**Fig. 5.**
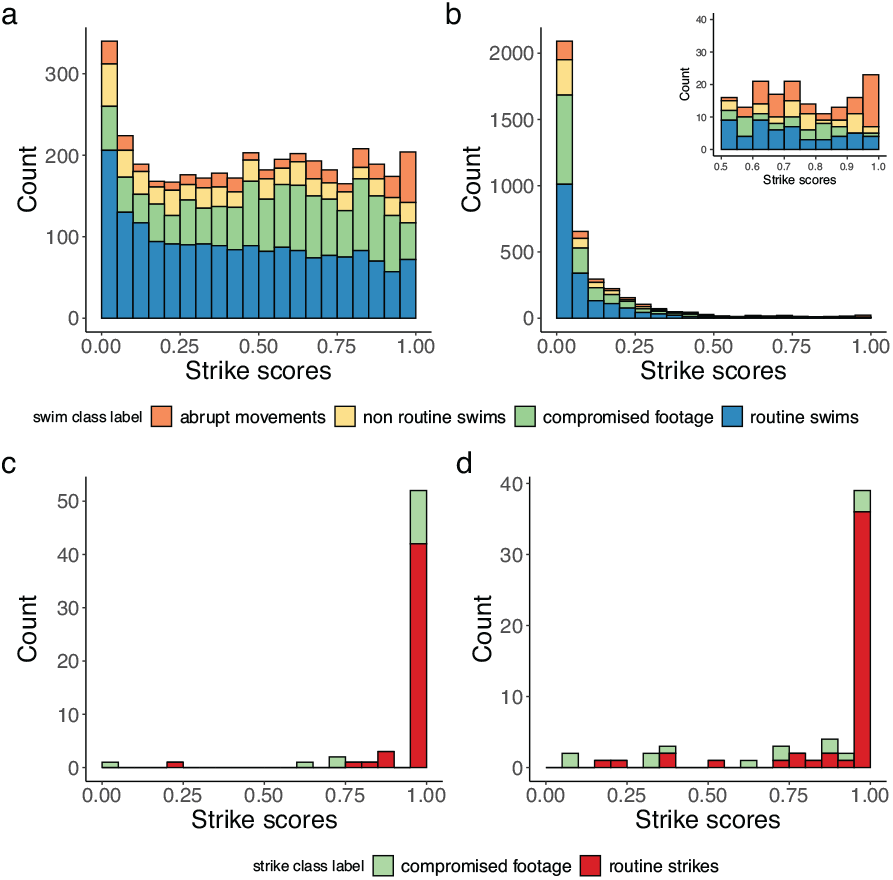
Error analysis. a,b) Distribution of strike scores and annotations for the naturalistic “swim” class as assigned by two SlowFast models a) Kinetics pre-trained, b) SSv2 pre-trained, inset is a close-up of the right tail of the distribution. Routine swimming appears in blue, compromised footage clips in green, non-routine swimming behavior (irregular, but not abrupt movements) in yellow, and abrupt movements (rapid, forceful movements) in orange. c,d) Distribution of strike scores for the naturalistic “strike” clips as assigned by the same models c) Kinetics pre-trained, d) SSv2 pre-trained. While SSv2 is better at classifying the “swim” clips, the Kinetics achieves superior results on the “strike” clips. Compromised footage and non-routine behaviors only partially explains mis-classifications.

We assessed the performance of our two classifier using only clips labelled as either high-quality routine swimming (n=1,841) or strikes (n=62). This new partition resulted in an AuROC of 0.95, 0.98 and AuPRC of 0.77, 0.84 for the Kinetics and SSv2 respectively. Note that this improvement in AuPRC can also be attributed to the change in class imbalance (∼ 3.4% versus the original ∼ 1.4%). To test whether model performance was adversely affected by one of the two filming setups, we further assessed the SSv2 variant separately on each of the filming setups (Fig. S6 in SI), results do not suggest any particular bias resulting from filming setup.

Given the low number of “strike” samples, stemming from the natural rarity of strike events in the video sequences, the cost of false negatives (missing strike events) is exceptionally high. We therefore figured that a human in the loop is needed for perfect classification. We used the strike score distributions of the two action classes (“swim”, “strike”) to calculate the expected number of clips that will need manual review under each of the variants (SSv2 and Kinetics; see SI section S5.4) when applying our pipeline to all 37 videos in the uncurated dataset. We estimate that this application will result in ∼ 870,000 clips. Our calculation reveal that the cost to recover 95% of strikes would be reviewing 156,600 clips (18%) for the SSv2 variant, and 252,300 clips (29%) for the Kinetics variant.

Considering the manual analysis of clips is much faster than that of entire videos, we estimate it at roughly 2.5 seconds per clip. Thus, analysis of the entire uncurated dataset, when aiming to obtain 95 % recall, using the current pipeline with the SSv2 variant will take 109 work hours. This is a substantial improvement over full manual analysis, amounting to approximately 20% of the original annotation effort.

## 5 Conclusions

In this study we have presented a data acquisition and analysis pipeline for the detection of larval feeding behavior in aquaculture rearing pools. We have created novel datasets featuring fish larvae freely behaving in unconstrained environments, under naturalistic conditions. On these datasets, we have benchmarked an action detection pipeline with several backbones in a curated manner. We show that training models on a smaller, highly curated dataset can yield good results also on a more difficult, naturalistic dataset, with a different structure of class imbalance. We found that even when pre-training data is from domains that are vastly different from monochrome underwater videos, pre-trained models achieve superior results.

Our work has shown that while it is feasible to use an analysis pipeline based on action classification for the detection of larval feeding, the different components of our system could be fine-tuned to eliminate those visual artifacts that impair the performance of our computer vision algorithms.

Our next two main algorithmic challenges lie in reducing false positive rates and moving to the uncurated realm. For the case of the Kinetics variant, in which routine “swims” are classified as “strikes”, techniques such as hard negative mining can prove beneficial. In the SSv2 case, hard positive mining is less practical as the frequency of the positive class in the raw data is extremely low. Given the extreme data imbalance and the huge annotation effort, we propose that an unsupervised approach based on anomaly detection, in which a model learns the “normal” class without any labels can be stronger in capturing events of interest. During our annotation process we encounter several videos without any strike events, these would be ideal for such a scheme.

For the uncurated challenge, we need to develop approaches that will enable the detection of strike behavior in long video sequences in realtime. A baseline approach would be to naively apply our pipeline with a sliding window approach throughout the video. Using approaches such as temporal localization of the actions is a logical next step, however it will require a huge annotation effort in order to annotate the temporal bounds of the tens of fish which appear in the videos.

Our work as presented here has established foundations for an *in-situ* underwater system for analyzing the feeding behavior of larval fish in aquaculture facilities. This system has the potential to reduce the knowledge gap in regard to the extreme larval mortality rate in the first weeks of their life; and will offer an effective monitoring and management tool for precision aquaculture.

## Supporting information

Video S1 (ESM_1)

Video S2 (ESM_2)

## Acknowledgments

We thank Keren Perry and Tamara Gurevich for their indispensable help with acquiring, annotating and analyzing larval feeding behavior manually. We also thank the entire Holzman lab for their help with the data labeling of the naturalistic dataset. We are indebted to Amir Jevnisek, Tal Perevolotsky and Anael Engel for inspiring brain-storming sessions. A special thanks to Moti Ohevia, Erez Levin, and the technical team of the Interuniversity Institute for Marine Science of Eilat for their help in the construction of the camera setups.

## Statements and declarations

The authors report no competing interests. This study was funded by the Israeli Ministry of Agriculture, Microsoft AI for Earth, and The Colton Family Next Generation Technological Institute and the Miles Nadal institute for Technological Entrepreneurship at Tel Aviv University.

All datasets described in this paper will become available upon acceptance through an appropriate data repository. Full code will be available in the paper’s Github repository, as well as trained model weights. The authors state that they comply with the ethical standards as defined by the journal.

## Author’s contribution

S.B: data curation, formal analysis, investigation, methodology, validation, visualization, writing—original draft and writing—review and editing; L.L: data acquisition, data curation, data analysis, methodology, writing—review and editing; S.A.: conceptualization, funding acquisition, methodology, resources, supervision and writing—original draft and writing—review and editing; R.H.: conceptualization, data curation, funding acquisition, methodology, resources, supervision, writing—original draft and writing—review and editing.

