## Supporting Information for "Analysis of Larval Fish Feeding Behavior under Naturalistic Conditions"

### Supplementary Information

In this supplementary information we provide the more in-depth and technical details of our work. We start with displaying an extended selection of samples from our dataset, including videos. We move on to describe in detail the training procedure of the various action classifiers we used; including an ablation study to test contribution of our variance image manipulation. This is followed by a description of our fish detection model and dataset. We then specify some of the implementation details for the curated experiment, followed by some additional results and error analysis. We present the labeling software used for annotating the naturalistic curated that resulted from this experimented. We end by providing a short explanation on our calculation of the expected performance of our two best models.

#### S1 Camera system examples

Over the course of 17 months, we filmed and annotated 37 videos using two distinct camera setups (see methods in main text). In Fig. S1 you can see some of the diversity in appearance that exists in our dataset. The figure shows frames extracted from the raw videos used to create our naturalistic dataset. Notably, see the difference in illumination, amount and type of prey in the water, number of fish, and various levels of sharpness of individual within and between frames. An example of some of the issues we had with each setup is also apparent, with most frames in the first setup (inside the dashed box) lacking crisp imagery, and some of the frames in the second setup showing the round edges of the housing (bottom frame column I and top frame column II).

Online Resource 1 is an example of a "swim" event from the naturalistic curated dataset, note the uneven lighting and multiple fish. Online Resource 2 is an example of a "strike" event from the balanced curated dataset. Note the even lighting typical of the balanced dataset and the characteristic S-shape and jump displayed by the fish.

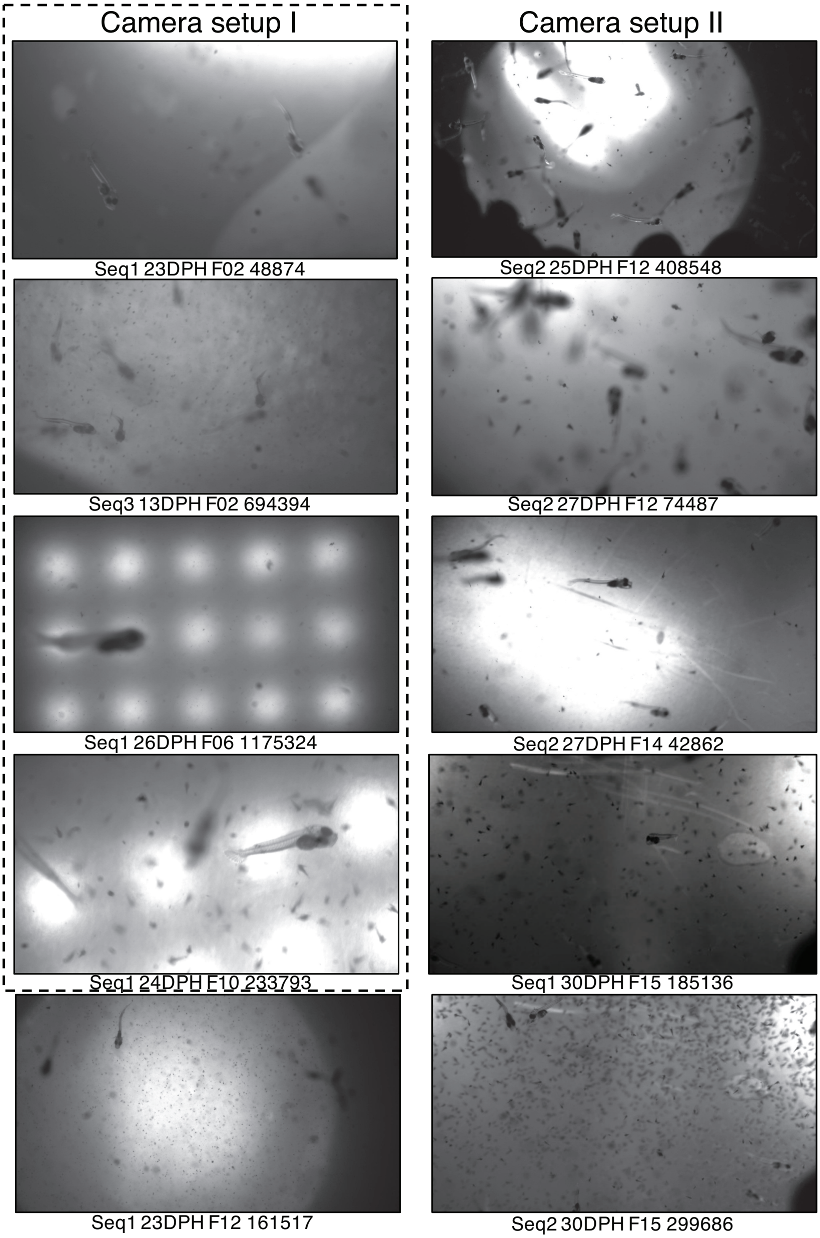


**Fig. S1**: Sample frames. Examples of video frames from various videos from the uncurated dataset, from both filming setups, the first in the dashed box, second outside it. Naming convention under each frame states the sequence number in that filming day (Seq); the age of the larvae (DPH, days post hatching); the cohort identity (F, each cohort was filmed consecutively throughout its growth cycle), and finally the frame number in the sequence. Different combinations of both technical and environmental variables create a visually diverse dataset

| Split | Swims | Strikes |
|---|---|---|
| Train | 41 | 39 |
| Validation | 11 | 11 |
| Test | 19 | 16 |

**Table S1**: Action classifier dataset composition.

### S2 Action Classifier training

For our action classifier, we trained all backbones (I3D, Slow, SlowFast) on our balanced curated dataset (see section 3.3.2 in the main text). I3D and Slow were pre-trained on Kinetics, SlowFast had three variants not pre-trained (None), Kinetics pre-trained, SomethingSomethingV2 (SSv2) pre-trained. We've split our data into the customary train/validation/test splits (see Table S1), making sure that clips from each video appeared in the same split. We further attempted to have roughly the same amount of clips from each of the two camera setups in each split. Even within each filming setup videos varied greatly in appearance due to changes in illumination, suspended particle density (food items) in the water, and fish density.

As mentioned in the main text, in our training, we used transfer learning, data augmentations and variance image manipulation to tackle our small sample size. in the next few paragraphs we'll expand on each of these topics. Additionally, we provide our full training and evaluation code, including all data augmentations, in the paper's Github repository.

#### S2.1 Training

We used the same protocol and hyper-parameters for all our experiments, both for testing different backbones and testing different pre-training strategies. For all pre-trained models we used pre-trained weights provided by the PyTorchVideo (Fan et al., 2021) library, replacing the classification head with a fully connected layer with 2,048 or 2,304 (Slow/I3D or SlowFast) inputs and 2 outputs; a strike neuron and a swim neuron. For the none-pretrained SlowFast network we simply initialized model weights at random using the default Pytorchvideo settings and replaced the classification head as described.

We fine-tuned the entire network on our Balanced dataset for 50 epochs using Stochastic Gradient Descent (SGD) optimizer with a learning rate of $1e-3$. We decreased learning rate throughout training using a ReduceLRonPlateau scheduler from PyTorch (Paszke et al., 2019), which reduces the learning rate if no improvement has been made in a chosen metric after a certain "patience" time. We set this metric to be the mean epoch training loss, and set the patience time to be 10 epochs.

We trained in batches of four 32-frame clips at a time (this was a constraint of the GPU memory). Typically in SlowFast, the user selects the number of crops per video and the number of clips, together defining the number of "views" per clip. We set this parameter to 1.0, because our behavioral events were extremely brief (dispensing the need to sample several clips), and our clips already cropped around single fish to ease training. To train the I3D network, we used an Azure Data Science Virtual Machine with an A100 NVIDIA GPU, as standard GPU memory was not sufficient for four clip batches of 32-frame clips. The Slow backbone took only the Slow pathway of the SlowFast network and was trained using batches of four 8-frame clips.

Seeing as our dataset is small, even a single mis-prediction by the classifier can shift the results. This is why we repeated the experiment using 3 different random seeds (16, 42, 82) and present the results averaged across seeds.

#### S2.2 Augmentations

During training we used augmentations to enhance our dataset. In our implementation we built upon the existing augmentations in the SlowFast and PyTorchVideo repositories. We also added two additional augmentations, brightness/intensity jitter and Gaussian blur, that suited the nature of our data.

We start by sub-sampling frames uniformly throughout the input clip (32 for the Fast path, 8 for the Slow path). We then applied random intensity jitter - a modulation of up to 20% (randomly selected) in intensity values, applied to the input at a probability of 0.5. This was followed by a random Gaussian blur, with a kernel of size (13,13) and a sigma of (6, 8). To apply Gaussian blur to the clip, we simply used TorchVision's (Paszke et al., 2019) Gaussian blur implementation and applied it to each frame in our clip. We then normalized the each channel in the video to have a mean pixel value of 0.45 with a standard deviation of 0.225, we tried removing this normalization in earlier trials but it gave better experimental results. Clips were then re-scaled to $256 \times 256$ pixel frames, to save on compute power. After this a set of flips and rotations was randomly applied at a probability of 0.5. These included random horizontal and vertical flip, and a random rotation by 90 degrees performed $k$ times, for some $k \in 1:4$, to get different rotations (90, 180, 270, 360 degrees).

These diverse augmentation meant to capture some of the diversity we saw in the fish that appeared in our dataset. From varying blurs, caused by fish moving in and out of the focal volume, to unconstrained 3D motion in all possible directions, caused by the fact we filmed in the water column of a large pool.

#### S2.3 Variance image implementation

We used what we call the variance image, to provide the network with additional movement-related information. We knew feeding strikes at prey were associated with extremely rapid movements, unlike swimming motion. As fish are dark blobs against a light background, we therefore hypothesized that the variance in the intensity of individual pixels in time will show different patterns during strike clips.

An example of variance images for each action class is given in Fig. S2. You can see that the variance image captured distinct movement patterns

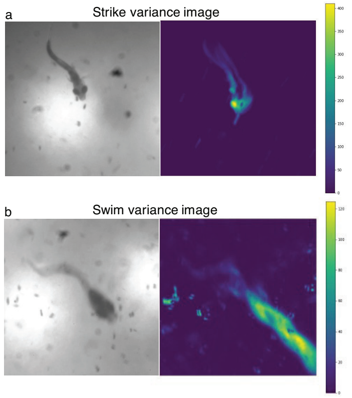


**Fig. S2**: Example of variance images. First frame in raw clip (left) and variance image on the entire clip (right) for a ”strike” clip (a) and a ”swim” clip (b). Note the different movement patterns captured in each clip by the variance image, and note the different scales on the right.

and variance amplitudes. The variance image, as its name implies, is simply a calculation of each pixel’s variance along the temporal axis of the clip. To amplify the effects the variance image captures, we applied a set of morphological operations using to create more diffuse areas with peak variance (see code). To insert the variance image calculation into our training, we chose to implement it as an additional augmentation. To insert the variance image with our input, we simply add it as additional channel with the unmanipulated monochrome frames; instead of an RGB we pass a 3-channeled input: (monochrome,monochrome,variance image).

#### S2.4 Results

After training, for 50 epochs we take the model from the last epoch of each experiment, and assess it on our validation and test sets. Results are averaged for each backbone across the three different random seeds.

As can be seen in Fig. S3 and Table S2, all pre-trained backbones classifier did well on the curated balanced dataset, reaching high levels of saturation in both the ROC and PRC, and correspondingly had high AuROCs and AuPRCs for all the dataset splits. However, clearly most SlowFast-based backbones achieve better performance than I3D, with the SSv2 pre-trained model showing the best results. The results of the Slow and SlowFast backbones pre-trained on Kinetics are comparable on this dataset, however diverge when assessed on the naturalistic dataset (see Fig 4 in main text). The performance of the SlowFast backbone with no pre-training is not better than a random classifier, exemplifying the importance of pre-training in our low-data regime. Due to the scarcity of data in the Balanced dataset we observed some apparent anomalies in our performance metrics (such as test performance being higher than validation).

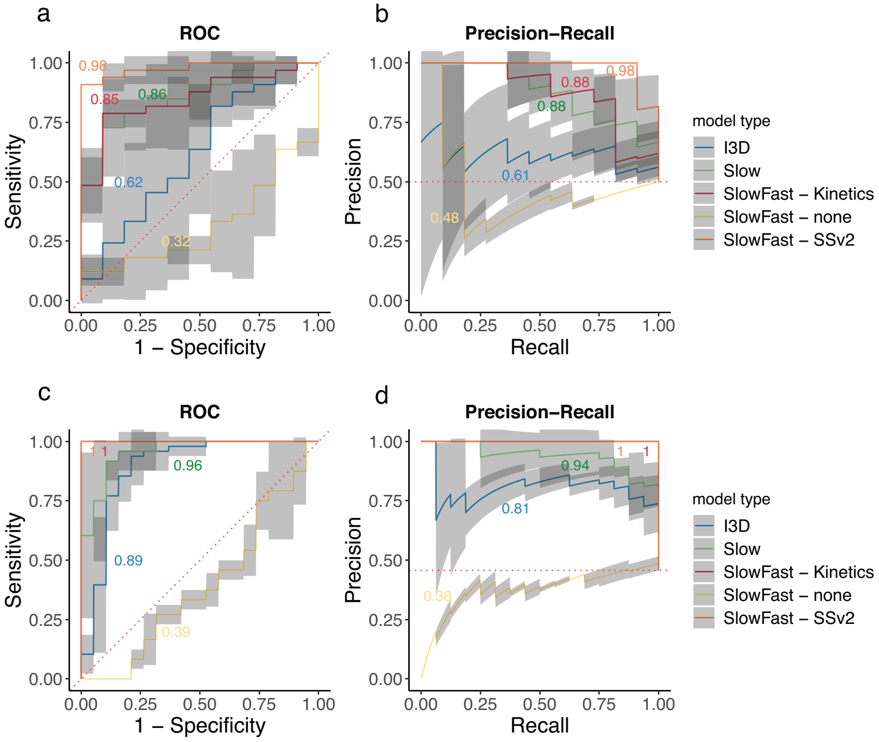


**Fig. S3**: Comparison of action classifier backbones and pre-training methods on the balanced curated dataset. The mean ROC and mean PRC for all models, averaged from training on three different random seeds. a,b) Results on the balanced validation set; c,d) Results for the balanced test set. Curves are presented for all backbones; I3D (blue), Slow (green), and SlowFast (red), all pre-trained on Kinetics; SlowFast with no pre-training (yellow) and SlowFast pre-trained on SSv2 (orange). The colored numbers are the area under each curve (AuROC/AuPRC). The shaded area around each curve represent the 0.95% confidence interval. The dashed red line represents the expected under a random untrained classifier. The pre-trained SlowFast backbones show superior performance.

##### S2.4.1 Variance image ablation study

To test the contribution of the variance image manipulation to our classifier training we trained SlowFast classifiers pre-trained on Kinetics with and without the manipulation. Seeing as our dataset is small, even a single misprediction by the classifier can greatly shift the results. This is why we repeated the experiment using 3 different random seeds (42, 17, 28), repeating each experiment twice per seed. This resulted in 12 trained models, 6 trained with variance image, and 6 without.

| Backbone | Train | | Val | | Test | |
|---|---|---|---|---|---|---|
| | AuROC | AuPRC | AuROC | AuPRC | AuROC | AuPRC |
| I3D | $0.99 \pm 0.014$ | $0.98 \pm 0.027$ | $0.62 \pm 0.147$ | $0.61 \pm 0.161$ | $0.89 \pm 0.036$ | $0.81 \pm 0.060$ |
| Slow | $1 \pm 0.003$ | $1 \pm 0.003$ | $0.86 \pm 0.093$ | $0.88 \pm 0.060$ | $0.96 \pm 0.028$ | $0.94 \pm 0.050$ |
| SlowFast - Kinetics | $1 \pm 0$ | $1 \pm 0$ | $0.85 \pm 0.043$ | $0.88 \pm 0.018$ | $1 \pm 0$ | $1 \pm 0$ |
| SlowFast - none | $0.63 \pm 0.008$ | $0.61 \pm 0.035$ | $0.32 \pm 0.0425$ | $0.48 \pm 0.017$ | $0.39 \pm 0.025$ | $0.38 \pm 0.007$ |
| SlowFast - SSv2 | $1 \pm 0$ | $1 \pm 0$ | $0.98 \pm 0.017$ | $0.98 \pm 0.011$ | $1 \pm 0$ | $1 \pm 0$ |

**Table S2**: Model performance on the balanced dataset. Numbers represent $meanAuC \pm std$ for each backbone, averaging results from training using three different random seeds, I3D and Slow are pre-trained on Kinetics, SlowFast backbones on Kinetics, SomethingSomethingV2 or with no pre-training.

We used the same training procedure described above, and after training, we selected the model from the epoch that performed best on the validation set. We plot in Fig. S4 the average ROC and PRC for all experiments on the validation set. The confidence intervals shown represent the 95% confidence bounds, calculated using the precrec package in R (Saito and Rehmsmeier, 2017).

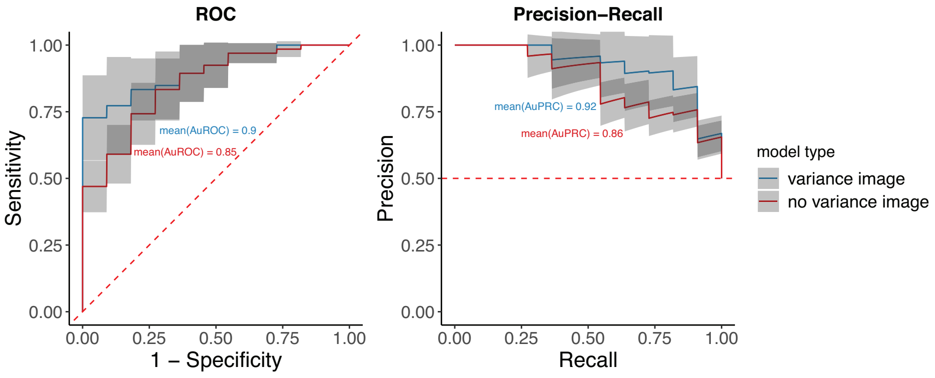


**Fig. S4**: Results for the ablation study on the efficacy of variance image manipulation. Mean ROC and PRC for 6 SlowFast models pre-trained on Kinetics trained with the variance image manipulation (blue), and 6 trained without the variance image manipulation (red). Confidence intervals are the 95% confidence bounds. Dashed red line is the expected under a random untrained classifier.

As can be seen in Fig. S4, while the average curves for the two conditions are not completely separated throughout their range, the use of variance image manipulation seems to afford some benefit in areas of low False Positive Rate (1-Specificity) in the ROC and areas of higher Recall in the PRC. In all experiments, the models trained with variance image were consistently higher scoring.

We expect that given enough data, our model should be able to learn such features regarding the movement of the fish on its own, without the "crutches" of image manipulation. However, as we don't have sufficient data as of yet, we show that this manipulation helped us achieve better results.

### S3 Fish detector dataset and training

Inspired by the SlowFast paper (Feichtenhofer et al., 2019), we also used the Detectron2 framework (Wu et al., 2019) to train an object classifier to detect our fish, so that we can extract clips centered around individuals from full-frame videos. Below we describe our dataset for training this detector, the training procedure and results.

#### S3.1 Dataset

Our fish detection dataset is composed of full-frames (1920 $\times$ 1080 pixels) extracted from videos and annotated with bounding boxes around each individual fish. We extracted frames from 4 different raw videos from our dataset, to cover a range of filming conditions.

From each video we sampled a 10 minute sequence, within which we further sampled 1 frame every 0.33 seconds to allow variability in fish number and positions. The selected frames were then annotated by research assistants, who fitted a tight bounding box around each fish in the frame.

In order to ease the labeling process, initial guesses for bounding boxes were generated using the same Canny edge detection-based methodology used in creating the balanced curated "swim" class (see section 3.3.2 in main text). These initial guesses were then fine-tuned by the research assistants and any missing fish added using the CVAT labeling tool (Sekachev et al., 2020). This process resulted in 1,664 annotated frames with 3,139 bounding boxes, split into 1,166 frames (2,196 boxes) in the train set, 245 frames (489 boxes) in the validation and 253 frames (454 boxes) in the test set. Unlike in the action classification dataset, we randomly assigned frames from all raw videos into the partitions. This was done in order to create a detector that will work well under various filming conditions. Except from one video, none of the raw videos in the detection dataset were used in the classification dataset. The video used in both appeared in the train partition of the classification dataset.

| Split | AP | AP50 |
|---|---|---|
| Train | 72.86 | 97.17 |
| Validation | 53.73 | 83.67 |
| Test | 50.39 | 85.05 |

**Table S3**: Fish detector training results for each split (first column), both average precision (AP, second column) and average precision at a threshold of 0.5 overlap with ground truth (AP50, third column), are shown

#### S3.2 Training procedure

Our detector was a Faster-RCNN (Ren et al., 2015) object detector with a ResNet-50-FPN backbone (He et al., 2016) (Lin et al., 2017). This detector was pre-trained on ImageNet (Deng et al., 2009) and fine-tuned on our detection dataset. We followed the recommended procedure in the official Detectron2 tutorial, for fine-tuning a pre-trained model on a custom dataset. We trained the model for 4,380 iterations, using 8 images per batch, a base learning rate of 0.00025, and, unlike the tutorial, we did not use warm-up in our training regime.

For the full set of hyperparameters and training procedure see the paper's repository.

#### S3.3 Detection training results

The overall average precision (AP) and average precision at a threshold of 0.5 overlap (AP50) for our Faster-R-CNN fish detector are recorded in Table S3, it can be seen that using a lower threshold yields far better results. Accordingly, we chose to set the bounding box score threshold at 0.5 for our pipeline.

### S4 Curated experiment implementation details

In this study we propose a pipeline for the detection of fish larvae feeding and test it in a curated experiment that resulted in the naturalistic curated dataset. Here we go into details regarding the method of sampling frames from the raw videos; the removal of overlapping clips; and the results on the naive set (see section 3.3.2 in main text).

#### S4.1 Frame sampling

For the experiment, We applied our pipeline to 11 raw videos known to contain the highest density of feeding strikes. For each video, we selected frames known to contain feeding strike or other interactions with prey, such as spitting; we specifically chose the mid-frame of each event (i.e., middle between start frame and end frame).

We additional sampled frames uniformly at random from the entire length of each video, keeping a ratio of ∼1:15 between frames that contained events of interest and those that did not. To prevent sampling too close temporally to the event of interest we did not sample ±25 frames around the mid-frame of each of the strike events. Frame numbers from which each clip is originated from are available as part of the naturalistic dataset's metadata.

#### S4.2 Overlapping clip removal

For frames containing many fish, it occurred that clips created based on the predictions of the fish detector were highly overlapping. To deal with this, we used a post-hoc heuristic to removed clips whose area was mostly overlapping, much in the spirit of non-maximal-suppression. Please see our code for full details.

This procedure resulted in the removal of 621 clips, roughly two thirds of which (391) from a single video, which had particularly dense fish distribution in the frames (Seq2 25DPH F12, top right column in Fig. S1). Our results are reported after this removal procedure.

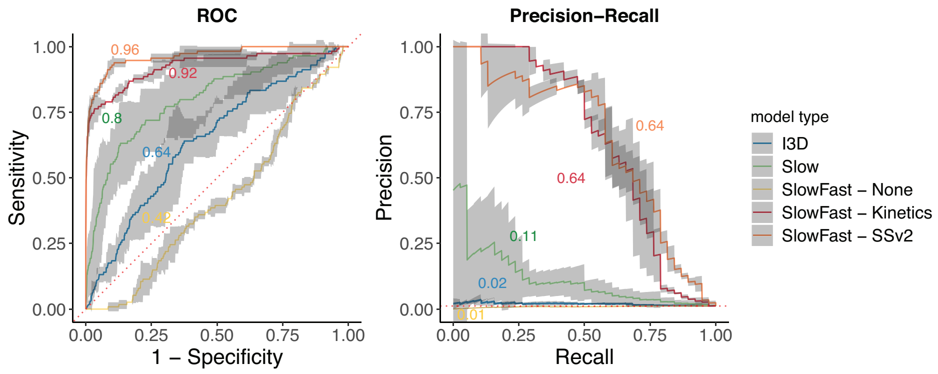


**Fig. S5**: Evaluation of backbones on the naive subset of the naturalistic curated dataset. a) mean ROC; b) mean PRC. Results are shown for Kinetics pre-trained I3D (blue), Kinetics pre-trained Slow (green),SlowFast with no pre-training (yellow), SlowFast pre-trained on Kinetics (red), SlowFast pre-trained on SSv2 (orange). Each curve is a mean of three models trained on different random seeds. Shaded area around each curve is the 95% confidence interval. In both a & b dashed red lines represent the performance expected by a random untrained classifier. Results show similar trends to those shown on the entire dataset. The SlowFast backbones are still the best-performing.

### S5 Curated experiment results and error analysis

#### S5.1 Naive set results

The naive set comprised clips from those raw videos that were not included in the train/validation splits of the balanced curated dataset (i.e., those video that were not source for clips the action classifier trained on). As can be seen in Fig. S5 these results show similar trends to those obtained on the entire naturalistic dataset with the pre-trained SlowFast models leading in both AuROC and AuPRC.

#### S5.2 Filming setup error analysis

We test whether one of the filming setup is introducing more error than the other, adversely affecting our pipeline's performance. To do this, we asses the performance of the SomethingSometingV2 (SSv2) pre-trained SlowFast models separately on the new and old filming setups.

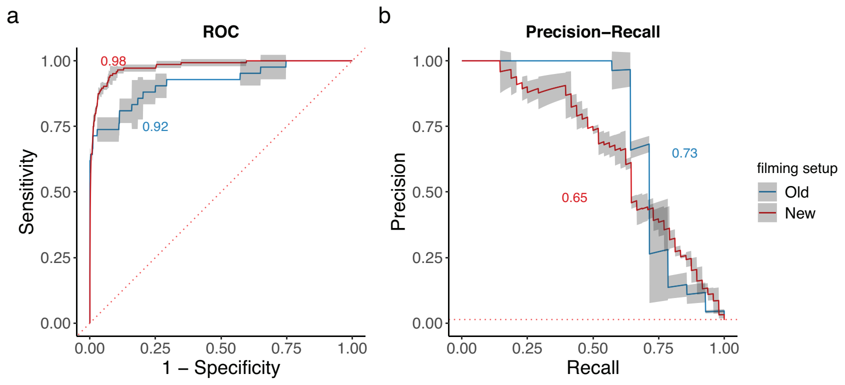


**Fig. S6**: Effect of filming setup on model performance. a) mean ROC, plotting the True Positive Rate (Recall/Sensitivity) against the of False Positive Rate (FPR/1-Specificity); b) mean PRC, presenting the Precision as a function of Recall. Results are shown for a SlowFast backbone pre-trained on SSv2 on the Old filming setup (blue) and New filming setup (red). Shaded area around each curve is the 95% confidence interval. In both a & b dashed red lines represent the performance expected by a random untrained classifier. The performance on the two filming setup appears comparable.

As can be seen in Fig SS6 there is an improvement in performance in the new filming setup in ROC, but not in PRC. This provides assurance that our improvement on the camera system does yield somewhat improved results, however this needs to be further examined, especially in light of the different class imbalances.

#### S5.3 Custom labeling software and details on naturalistic dataset annotations

After concluding the curated experiment, we were left with 4,563 clips, 62 of which we knew contained strike events, but we did not know much about the rest.

We were particularly interested in the apparently high rate of false positives we were seeing. Preliminary exploration led us to believe there were some consistent biases or reasons behind these mistakes. So we recruited the help of unbiased observers to which we gave many possible labels to choose from, which are depicted in Fig. S7.

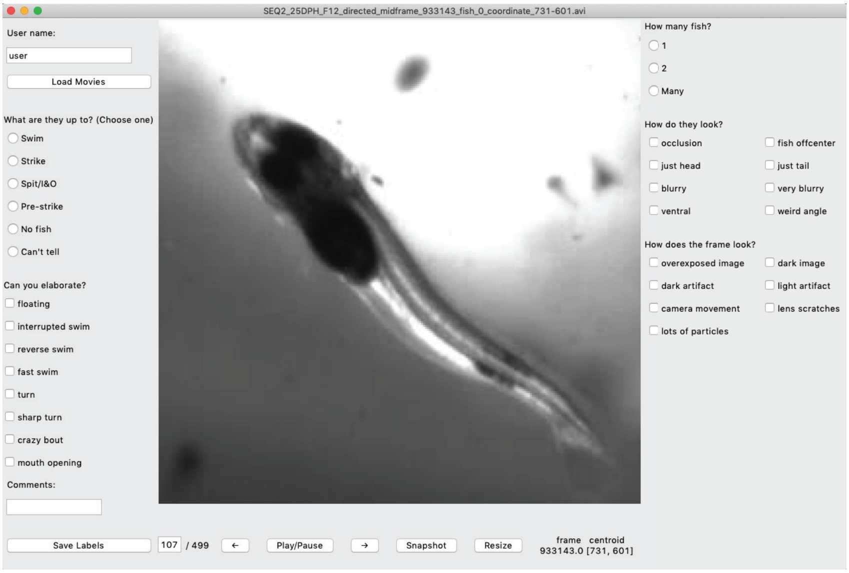


**Fig. S7**: Custom Labeling Software. Developed using python3, Tkinter, and OpenCV. Main activity on the top left column is the main behavior exhibited by the fish, below this are some more finer grain details about the motion of the fish. To the right of the video, parameters about the amount of fish appearing in the frame, their appearance (including whether they are occluded) and the general appearance of the frame are collected. This allowed our labelers to create a rich description of the behavioral and visual parameters displayed in the clip.

We collected information about the behavior of the fish; both broadly (top left radio buttons) and more fine grained motions (bottom left, check buttons). We also collected information about the number of fish visible, the appearance of the fish themselves and the appearance of the frame in general (right side of the panel).

We later merged these into the four main categories discussed in the main text: strikes, abrupt movements, non-routine swimming, compromised footage, routine swimming, and can't tell or No fish.

Abrupt movements refer to clips in which the larvae rapidly changed swimming direction or attempted to spit out prey items. Non-routine swimming refers to clips showing floating (no undulations of the body or fins), interrupted swimming (rapidly accelerating or decelerating), and reverse swimming. Compromised footage refers to overexposed images, caused by the filming setup; fish appearing to move unreasonably fast in/out of frame as the result of strong local flows; and very blurry footage caused by fish being outside the focal volume. "Can't tell" category was typically used in cases in which the focal fish was occluded or the image was too dark to describe its behavior. "No fish" refers to cases of false identification by the detection module. Strikes were defined as rapid lunges towards the prey followed by opening of the larva's mouth. All other samples were considered routine swimming.

#### S5.4 Performance analysis - calculating expected workload for a human in the loop

We investigated the expected performance of our two top performing models - Kinetics pre-trained SlowFast and SSv2 pre-trained SlowFast, by calculating the projected number of clips we'll have to review to recall 95% of the strike events in our data (see section 4.2 in main text). To calculate this, we look at our two score distributions - for the "swim" and "strike" classes. The clips we will have to go over are the True Positives and the False Positives - all those clips that score higher than some decision threshold $x$ between $[0, 1]$. We run a cumulative sums on the counts of each of the score histograms (from highest to lowest scores), for the "swim" and "strike" classes. This sum will give us the number of clips we need to review up to each score bin/threshold in the histogram. We combine the two sums to get the total number of clips we will have to review, divided by the sum of all clips in both distribution we get the percent of clips to be reviewed. We then plot this number as a function of recall (i.e., the cumulative sum for the counts of the "strike" class divided by the total of strikes) in Fig. S8.

As can be seen, the SSv2 pre-trained model is better throughout, however the gap in performance diminishes near the extremes of the plot (very low/high recall). To get 97 % recall, the Kinetics model will have a human analyze roughly 40% of the videos generated by the system, while the SSv2 will do the same work for around 31 %. It is true the Kinetics doesn't get very low recall, due to the good mapping of the "strike" class to high strike scores, but the price of False Positives is still so high, that at whatever point you choose on the graph, the SSv2 variant will yield less clips for the same recall.

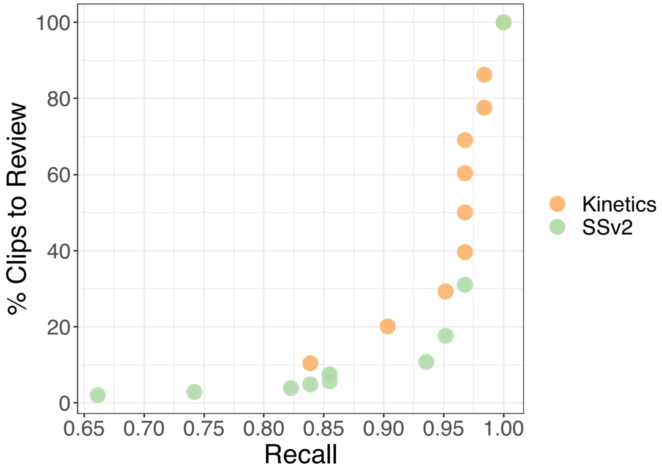


**Fig. S8**: Expected workload for a human in the loop. The percent of clips to review (true positives and false positives) as a function of recall. The Kinetics pre-trained model (orange) consistently yields a higher amount of clips to analyze, compared to SSv2 (green). However, the price of 100 % recall is the same for both models.
